# Genotyping Short Tandem Repeats Across Copy Number Alterations, Aneuploidies, and Polyploid Organisms

**DOI:** 10.1101/2024.12.13.628141

**Authors:** Max A. Verbiest, Elena Grassi, Andrea Bertotti, Maria Anisimova

## Abstract

Short tandem repeats (STRs) are a rich source of genetic variation, but are difficult to genotype. While specialized repeat variant callers exist, they typically assume a euploid human genome. This means recent findings regarding phenotypic effects of STR variants in human health and disease cannot be readily extended to polyploid organisms or cancer, which is characterised by copy number alterations (CNAs). Here we present ConSTRain, a novel STR variant caller that explicitly accounts for the copy number of loci in its genotyping approach. We benchmark ConSTRain using a euploid human 100X whole genome sequencing sample where it calls STR allele lengths for over 1.7 *×* 10^6^ loci in under 20 minutes with an accuracy of 98.28%. Subsequently, we show that ConSTRain resolves complex STR genotypes in an artificial trisomy 21 sample and a polyploid Dwarf Cavendish banana harbouring a large duplication. Finally, we analyse a microsatellite instable colorectal cancer tumoroid, where ConSTRain tackles CNAs and whole-genome duplications. ConSTRain is the first STR variant caller that allows for the investigation of repeats affected by CNAs, aneuploidies, and polyploid genomes. This unlocks the investigation of STRs across a wide range of contexts and organisms where they previously could not be easily studied.

## Introduction

Short tandem repeats (STRs), also known as microsatellites, are genomic regions where a DNA motif one to six base pairs (bp) in length is repeated consecutively. STRs are highly variable. Especially prevalent are insertion and deletion (indel) mutations that expand or contract the repeat by one or more unit [1]. Such STR variants may cause frameshift mutations or affect the phenotype by regulating gene expression levels in health and disease [2, 3, 4]. STR loci for which the allele length is associated with gene expression levels are called expression STRs (eSTRs).

The distinct mutational characteristics of STRs cause issues when genotyping them with general-purpose variant calling tools. To this end, specialized STR variant calling algorithms have been developed [5, 6, 7, 8]. While these tools enable accurate variant calling of STR loci from human sequencing samples, there are several key points they do not address. Notably, current STR genotypers were developed with the euploid human genome in mind. This means such tools expect two copies of each repeat locus to be present, with some tools supporting a ploidy of one for sex chromosomes.

While this may generally hold for mammalian genomes, it is not representative of the full range of genomic variation. Copy number alterations (CNAs) can change the ploidy of parts of a chromosome — and thus of the STRs located in those regions. CNAs can be present in the germline of healthy individuals [9]. Furthermore, somatic CNAs are a key feature of cancer, where they contribute to carcinogenesis by deleting and upregulating biological functions [10]. There are also more extreme cases where the ploidies of whole chromosomes (e.g., trisomy 21) or the full genome (i.e., whole-genome duplications) are affected. We recently described a panel of putative eSTRs in colorectal cancer [4]. However, since current STR variant callers do not account for CNAs, our eSTR detection approach had to exclude all STRs that were located in regions affected by CNAs. This lead to a substantial fraction of information — around 15% of all calls — being removed, meaning we may have missed important eSTR loci. Besides not addressing aneuploidies or CNAs, the focus of current STR variant callers on the human genome also means that such tools cannot be readily used to study STRs in polyploid organisms. While polyploidy occurs sporadically in animals, it is widespread in plants [11]. Among the polyploid plants are many important food crops like wheat, maize, and banana [11, 12, 13]. Despite the societal importance of such species, current computational tools do not allow for the extension of findings regarding the phenotypic effects of STR variants to polyploid organisms.

To address these open issues, we here introduce a new STR variant caller named ConSTRain (**<u>co</u>**py **<u>n</u>**umber guided **<u>STR</u> <u>a</u>**llele **<u>in</u>**ference). The fundamental idea of ConSTRain is that the copy number of each STR locus is explicitly considered in the variant calling process. The copy number can be set at the chromosome level by specifying the karyotype of the organism. Furthermore, ConSTRain allows the copy numbers of specific genomic regions to be changed by specifying CNAs known to be present in a sample.

We demonstrate that our new method is highly competitive: ConSTRain’s accuracy is at least as high as state-of-the-art STR variant callers on a euploid human benchmark, while the runtime is substantially lower (especially when running multithreaded). Furthermore, we apply ConSTRain in aneuploid settings on simulated trisomy 21 data, and on whole genome sequencing (WGS) data from a triploid *Musa acuminata* Dwarf Cavendish banana. The original publication of this *M. acuminata* sequencing data reported a large duplication on the long arm of chromosome 2 [13]. We show that ConSTRain is able to account for this duplication when the coordinates of the affected region are provided. Finally, we analyse STRs in four WGS samples from a microsatellite instable (MSI) colorectal cancer (CRC) tumoroid [14, 15]. One of these samples represents the original tumoroid line and the other three are clonal organoids, two of which have undergone whole-genome duplication. While these samples stem from the same tumour, we observe differences in STR allele lengths in pairwise sample comparisons. This indicates that ConSTRain can be useful for analysing tumour heterogeneity and tracing clonal lineages in cancer, even in closely related samples. Overall, ConSTRain is a flexible, fast, and accurate STR variant caller that can genotype repeats in human and non-human sequencing data while addressing ploidy-altering events.

## Materials and Methods

### ConSTRain implementation

ConSTRain is an STR variant caller implemented in Rust. It relies on the htslib C library [16] through the rust-htslib crate [17]. All analyses reported in this manuscript were performed using ConSTRain version 0.9.1. A visual overview of ConSTRain’s genotyping approach is shown in Figure 1. ConSTRain requires three input files: an alignment of sequencing reads to a reference genome (SAM/BAM/CRAM format), a file specifying the locations of STR loci (BED format), and a file specifying the karyotype (i.e, the ploidy of each chromosome) (JSON format). If the alignment file is in CRAM format, the reference genome must also be supplied (FASTA format). Optionally, a file specifying the location and copy number of regions affected by CNAs can be supplied (BED format). The estimated STR genotypes for each locus in the input STR panel are written to stdout in VCF format. Source code, details of input and output file formats, and an overview of available command line arguments are available at https://github.com/acg-team/ConSTRain.

**Fig. 1:**
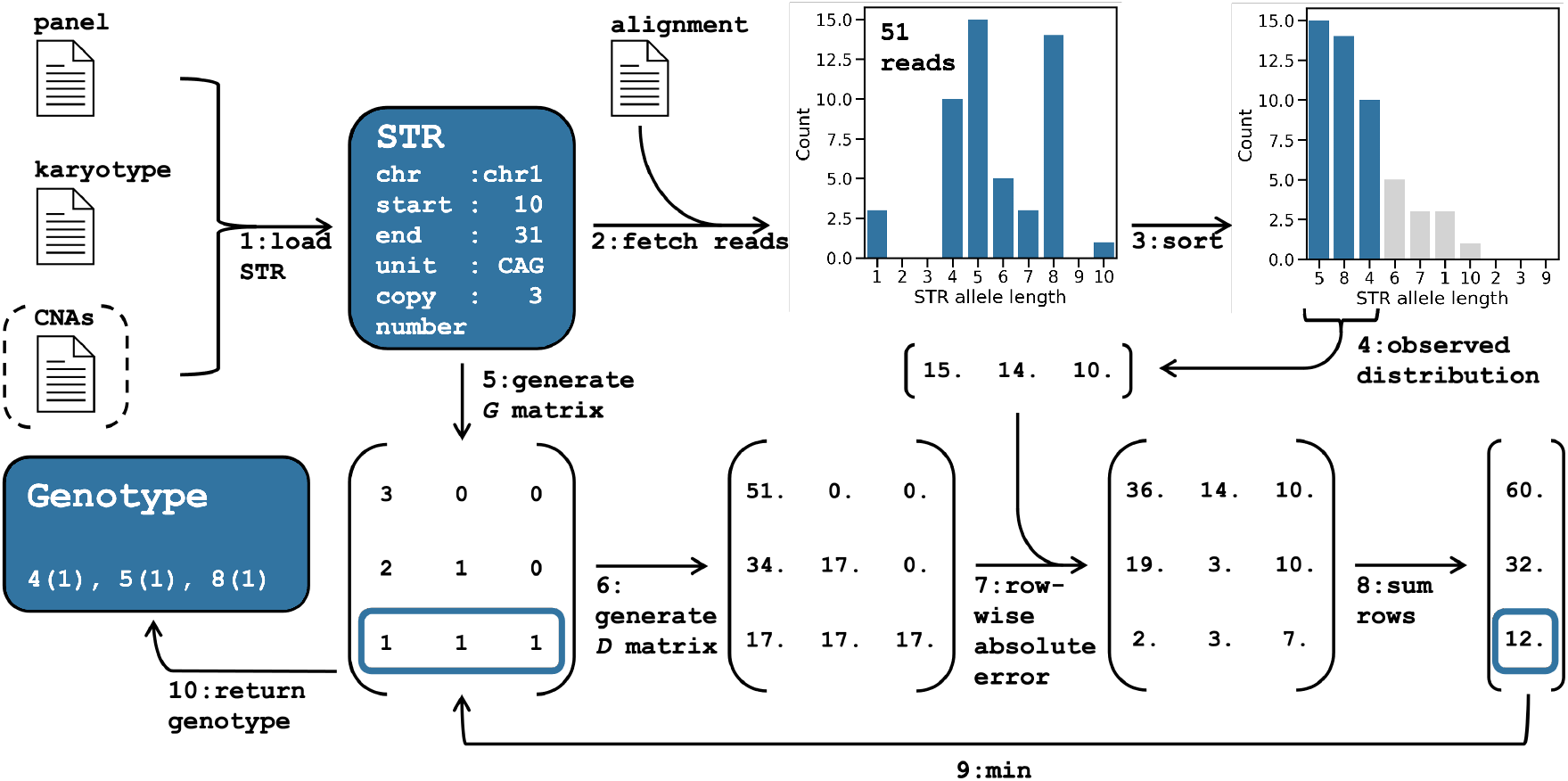
ConSTRain overview and example. **(1)** An STR locus are loaded from the input files. The locus reference information is parsed from the STR panel. The STR copy number is set based on the karyotype, and optionally updated if the STR is affected by a CNA. **(2)** Reads overlapping the STR region are extracted from the alignment file, and the length of the STR region in each read is determined. **(3)** The observed distribution is sorted, and at most as many allele lengths as the STR copy number are kept. **(4)** This yields the final observed allele length distribution. **(5)** Next, all possible genotypes are generated for the STR copy number and stored in matrix ***G*. (6)** From ***G***, the matrix ***D*** is generated by multiplying it with the total number of mapped reads (51 in the example) divided by the STR copy number (3 in the example). Each row in ***D*** corresponds to the expected allele length distribution of one of the genotypes in ***G*. (7)** The expected distribution with the lowest error to the observed distribution is found by taking the absolute difference between each row in ***D*** and the observed distribution, then **(8)** taking the sum of rows and finding the one with the lowest value. **(9)** The genotype in ***G*** with the lowest error is selected **(10)** and reported in the output. The inferred genotype of the STR locus in this example consists of an allele of 4 CAG units (present once), an allele of 5 CAG units (present once), and an allele of 8 CAG units (also present once).

### Vocabulary

Different fields and research groups use inconsistent terminology to describe the various characteristics of STRs. To avoid confusion, we will explicitly define the vocabulary used by ConSTRain here: STRs are made up of a sequence of repeated *units*. Currently, ConSTRain allows only perfect STRs, without any mismatches, insertions, or deletions between the different units of a locus. The number of nucleotides in the unit is referred to as the *period*. The number of times a unit is repeated is called the STR *allele length*. A *genotype* is the combination of allele lengths that exist for an STR locus in a sample. Finally, each STR has a *copy number*, which indicates how many instances of the STR locus exist in the genome.

### Initialisation

ConSTRain starts by reading STR loci from the STR panel file and the ploidy of contigs from the karyotype file. The copy number of an STR is initially set based on the ploidy of the contig it is located on. E.g., the copy number will typically be set to two for STRs located on human autosomal chromosomes. However, if a file with CNAs is provided and the coordinates of the STR intersect with the coordinates of a CNA, the copy number of the STR is updated to that of the CNA. Subsequently, ConSTRain fetches all reads from the alignment file that fully span the STR locus and parses CIGAR strings to extract the STR allele length from each read. This yields the observed allele length distribution for that STR. For reasons that are discussed below in ‘Generating all possible genotypes’, the distribution is sorted such that the STR allele lengths are listed according to their observed frequencies, in descending order (Figure 1, step 3). The main task for ConSTRain is to infer the most likely genotype for each STR locus, given its observed allele length distribution and copy number.

### Estimating the most likely STR genotype

Rather than using a heuristic optimisation approach to estimate the most likely genotype, ConSTRain explicitly generates all possible genotypes for an STR locus. From each of these possible genotypes, an expected allele length distribution is generated. The genotype for which the expected allele length distribution has the lowest absolute error (Manhattan distance) to the observed allele length distribution is chosen as the most likely genotype. To make this process tractable, ConSTRain operates under three assumptions:

1. STRs exist at integer copy numbers.
2. There are at most as many distinct allele lengths as the STR copy number.
3. Each STR allele in the genotype contributes an equal number of reads to the allele length distribution.

Under these assumptions ConSTRain can generate all possible genotypes for an STR locus, given its copy number. ConSTRain only considers genotypes where the alleles are in descending order of abundance (‘Generating all possible genotypes’ for details). Thus, if the STR has copy number two the possible genotypes are ‘AA’ and ‘AB’, if the copy number is three, the possible genotypes are ‘AAA’, ‘AAB’, and ‘ABC’, etc. Internally, ConSTRain represents genotypes as matrices. E.g., for copy number three the possible genotypes are:

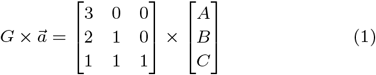

where each row in G represents a possible genotype, each column represents an STR allele length (encoded in 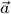), and each value represents the number of times an STR allele length is present in a genotype. For a given locus, the shape of G will always be such that the number of columns equals the copy number of the locus, and the number of rows equals the number of integer partitions that exist for that copy number (‘Generating all possible genotypes’ for details). Next, ConSTRain uses G to generate a matrix of expected allele length distributions, denoted as D. To do this, ConSTRain must first know the number of reads each allele in the genotype is expected to contribute to the STR allele length distribution. Under assumptions (2) and (3) we can find this number by dividing the total number of reads mapped to the STR locus by the copy number of the STR locus. ConSTRain multiplies G by this scalar which results in matrix D where each row contains the expected allele length distribution for the corresponding row in G. For each row in D, the absolute error to the observed allele length distribution of the STR is calculated. If the number of allele lengths observed for a locus is greater than the copy number (such as in Figure 1) only as many alleles as the copy number are considered. Conversely, if the number of observed alleles is fewer than the copy number, zero values are appended to the observed allele length distribution until its length equals the copy number. ConSTRain reports the genotype in G for which the associated expected allele length distribution in D has the lowest error to the observed allele length distribution. In cases where multiple genotypes are equally likely, ConSTRain does not report a genotype and sets a VCF filter tag to indicate why a genotype was not inferred. However, the observed allele length distribution will still be included in the VCF record’s FORMAT field.

### Generating all possible genotypes

As noted above, ConSTRain does not consider genotypes where allele abundances are not in descending order. Under assumption (3) it is not possible for an STR allele that is less abundant to contribute more reads to the allele length distribution than another allele that is more abundant. By sorting the observed allele length distribution we thus do not need to consider genotypes with non-descending allele abundances. Genotypes of this form would result in expected allele length distributions that are impossible under ConSTRain’s assumptions. Sorting the observed allele length distribution is a way to reduce the combinatorial space of possible genotypes: without doing this the number of genotypes to generate for a locus would be equal to the number of weak integer compositions of a size equal to the STR copy number. A weak integer composition refers to the representation of an integer as the sum of a sequence of non-negative integers. For a given integer, the number of weak compositions of a specific size (i.e., the number of terms to represent the integer as) is given by:

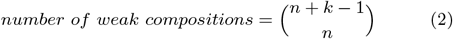

where n is the integer and k is the composition size. For our purposes n equals k equals the STR copy number. By sorting the observed allele length distribution we instead only need to generate a number of genotypes equal to the number of integer partitions of the STR copy number. Integer partitions differ from compositions in that the terms of the partition are not ordered, i.e., different orders of the same terms are considered identical. No closed-form solution is known to determine the number of partitions for an integer, but Sloane’s sequence A000041 enumerates the number of partitions for a range of integers [18]. Going back to the example in equation Equation 1, we can use Equation 2 to calculate that there exist ten weak integer compositions when n and k are both three. On the other hand, A000041 tells us that there are three integer partitions — a difference of seven. This may not seem very impactful, but the difference becomes much more pronounced for higher copy numbers: for n = 20 there exist more than 68.9 *×* 10^9^ weak compositions of size 20, but only 627 partitions. Further, given that an STR panel can contain hundreds of thousands of loci (e.g., over 1.7 *×* 10^6^ for the human genome), even small optimisations make a difference in overall runtime.

### Updating an existing VCF file

Besides the standard mode of running ConSTRain outlined above, ConSTRain also supports reanalysing previously generated VCF files. This may be useful if novel CNA information for a sample becomes available after an input alignment has already been analysed, or if it is necessary to adjust filtering parameters. It also prevents having to re-download large alignment files from remote repositories. This is possible because ConSTRain includes the observed allele length distribution of each STR in a FORMAT field of the output VCF. Since it is much faster to read the observed allele length distribution from a VCF file than to extract it from sequencing reads in an alignment, running ConSTRain in this mode is typically a matter of seconds.

### Filtering ConSTRain output

Genomic regions where the depth of coverage is lower or higher than expected may indicate a large number of technical artifacts for that region. This can lead to inaccurate variant calls. To address this, ConSTRain allows for the filtering of STR loci based on their normalised depth of coverage. The normalised depth is calculated by dividing the number of mapped reads by the locus copy number. This normalisation is important because the copy number of a locus is expected to affect the depth of coverage. When analysing an alignment of human male sequencing reads, for instance, loci on the sex chromosomes are expected to have roughly half the depth of coverage as loci on autosomes. Similar effects exist for genomic regions that are amplified or deleted by structural variants. Dividing the depth of coverage by the locus copy number will force all loci to occupy the same range of normalised depth values, which makes filtering more straightforward. The desired minimum and maximum normalised depth values can be set at the command line via the `--min-norm-depth` (default: 1.0) and `--max-norm-depth` (not set by default) arguments, respectively. These upper and lower bounds can be set manually to reasonable values before running ConSTRain. Another option is to first run ConSTRain without filters and then set bounds based on the observed distribution of normalised sequencing depths across all loci in the sample. This can help identify the range of acceptable normalised depth values for a specific sample. Once the minimum and maximum values are found, ConSTRain can be rerun on the VCF file with the updated filtering parameters. Since running ConSTRain on a VCF file is extremely fast (around 20 seconds for 1733646 loci on a 2020 MacBook Pro) this only marginally increases the overall computational workload. A Python script to generate a distribution of normalised depth values from a ConSTRain VCF file is included in the ConSTRain GitHub repository.

### STR reference panels

ConSTRain needs a reference panel of STR loci to know where STRs are located in the reference genome. The reference panel that was used in all experiments involving human data reported in this manuscript is based on the GRCh38 version 13 reference panel provided by GangSTR [7]. While ConSTRain is primarily aimed at genotyping STRs with periods between one and six, the repeat panel provided by GangSTR contains a small number (20481) of repeat loci with longer periods (up to 20), which we did not remove. Furthermore, the GangSTR panel does not contain mononucleotide repeats. We therefore extended this panel to include perfect mononucleotide repeats of at least allele length ten, which were identified using mreps [19]. This resulted in a panel containing 1733646 repeat loci in the GRCh38 human reference genome (Supplementary Figure 1A). The total region length of most of these repeats was relatively short, with only 3.38% being longer than 30bp in the reference assembly (Supplementary Figure 1B). This suggests that — barring large expansions — the vast majority of repeat loci in this panel should be resolvable with short sequencing reads.

We created a novel STR referene panel for the DH-Pahang v4 banana reference genome [20]. This was also done using mreps [19], setting command line arguments such that perfect repeats with periods one through six were reported. We subsequently filtered mreps output using custom Python scripts to retain only perfect STRs with at least allele length ten, six, four, three, three, and three for STRs with period one through six, respectively. This yielded a reference panel of 183345 STR loci across the 11 main chromosomes in the DH-Pahang v4 reference.

### HG002 benchmark

STR genotyping tools were benchmarked using haplotypes provided by the telomere-to-telomere (T2T) consortium’s Q100 project [21, 22]. The Q100 project provides high-quality, phased haplotypes of the HG002 cell line which have been used previously to benchmark STR variant calls [22]. To obtain ground-truth allele lengths for loci in our STR reference panel in the HG002 cell line, the Q100 haplotypes were mapped to the GRCh38 reference genome using minimap2 [23]. The resulting PAF file was parsed to find a ground-truth STR allele lengths in GRCh38 coordinate space. The HG002 allele length could be recovered for 1695865 STR loci in our panel (97.82% of the total).

Subsequently, STR variant callers were used to genotype the STR reference panel in an alignment of 2×250 Illumina whole-genome sequencing reads of HG002, which is available through Genome in a Bottle [24]. STR allele lengths generated by the different variant callers were compared to the ground-truth allele lengths derived from the Q100 haplotypes. Genotyping accuracy was calculated by determining the fraction of loci for which the biallelic genotypes reported by a variant caller exactly matched the allele lengths observed in the Q100 haplotypes.

### Simulating trisomy 21

We simulated 2×150 paired-end reads from chromosome 21 of the maternal and paternal haplotypes of HG002, as well as GRCh38. Since we were not interested in modelling sequencing errors, we simulated error-free, paired-end reads to a depth of coverage of 15X for each of the three haplotypes using wgsim (https://github.com/lh3/wgsim). Simulated reads from the three haplotypes were then combined to form a 45X sequencing sample of a triploid chromosome 21. These reads were mapped back to the GRCh38 reference genome using minimap2 [23].

### *Musa acuminata* whole-genome sequencing data

The *M. acuminata* sequencing data used here consist of two sequencing experiments of the same organism, one performed on an Illumina HiSeq1500 machine and the other on an Illumina NextSeq500 [13]. We downloaded all sequencing reads (European Nucleotide Archive, study PRJEB33317) and combined outputs of sequencing runs into two FASTQ files, one for the HiSeq1500, one for the NextSeq500. The two FASTQ files were mapped to the DH-Pahang v4 reference genome [20] using minimap2 [23], removing improper pairs, duplicate alignments, and low-quality alignments. These alignments will be referred to as the ‘HiSeq1500 alignment’ and the ‘NextSeq500 alignment’. Subsequently, the HiSeq1500 and NextSeq500 alignments were concatenated to form the ‘merged alignment’.

### Colorectal cancer whole-genome sequencing data

We obtained WGS data of a patient-derived cancer CRC tumoroid generated as a part of a previously published mutation accumulation experiment [14]. These data are available through the European Genome-phenome Archive under accession number EGAD50000000411. Briefly, this experiment was set up so that individual cells were isolated from a CRC tumoroid [25] and allowed to grow for six weeks. At the six week mark, WGS was performed on each clone to obtain a high-quality representation of the genome of the individually isolated cells. Subsequently, clones were repeatedly bottlenecked to 100 cells every two weeks for six months, followed by WGS of the resulting clones [14].

As a demonstration of ConSTRain’s applicability to cancer sequencing data we analysed four WGS samples from a single microsatellite instable tumoroid. The first of these samples was taken from the original tumoroid line and the other samples represented three different clones (01-0, 05-0, and 07-0) sequenced after six weeks of growth. For each sample, CNA calls generated by Sequenza were available [14, 26]. These CNA calls indicated that while the original tumoroid line and the 05-0 clone were diploid, the 01-0 and 07-0 clones had undergone whole-genome duplications and were tetraploid. We ran ConSTRain on all four samples, providing the appropriate Sequenza CNA calls each time. Then, we computed pairwise STR-based distances between samples based on the genotypes returned by ConSTRain. We limited this analysis to high confidence STR genotypes where the unit size was between three and six and the normalised depth of coverage was at least 5. For comparisons between diploid and tetraploid samples all genotypes in the diploid sample were artificially duplicated before performing comparisons. This meant that the diploid genotype [10, 10] would be represented as [10, 10, 10, 10], and [8, 9] as [8, 8, 9, 9], etc. Loci that were annotated with a different copy number in the two samples of a pair were not considered when calculating pairwise distances. The impact of this filter varied depending on which two samples were being compared: when comparing the two 2n samples 4.81% of loci did not have the same copy number, whereas up to 26.85% of loci had to be removed when comparing 4n samples. This is likely due to the fact that accurately calling copy number levels from sequencing data is a difficult task, especially for higher copy numbers. Pairwise sample distances were calculated by taking the sum of Manhattan distances between STR genotypes for all loci with a high confidence call in both samples, normalising by the total number of compared loci. This resulted in a distance between samples with a unit of ‘average difference in allele length per locus’.

## Results

### ConSTRain accurately genotypes STRs in euploid human sequencing data

We first evaluated ConSTRain’s performance when analysing sequencing data from a euploid human genome. We ran ConSTRain with default parameters on 100X short-read WGS data of the HG002 human cell line. Using high-quality HG002 assemblies as ground truth, we determined ConSTRain’s accuracy (Figure 2A&B, Supplementary Figure 3). ConSTRain returned allele length estimates for 1655655 out of the 1695865 repeat loci (97.63%) for which a ground truth was available. For 95.25% of these, the allele length(s) returned by ConSTRain exactly matched those of the ground truth.

**Fig. 2:**
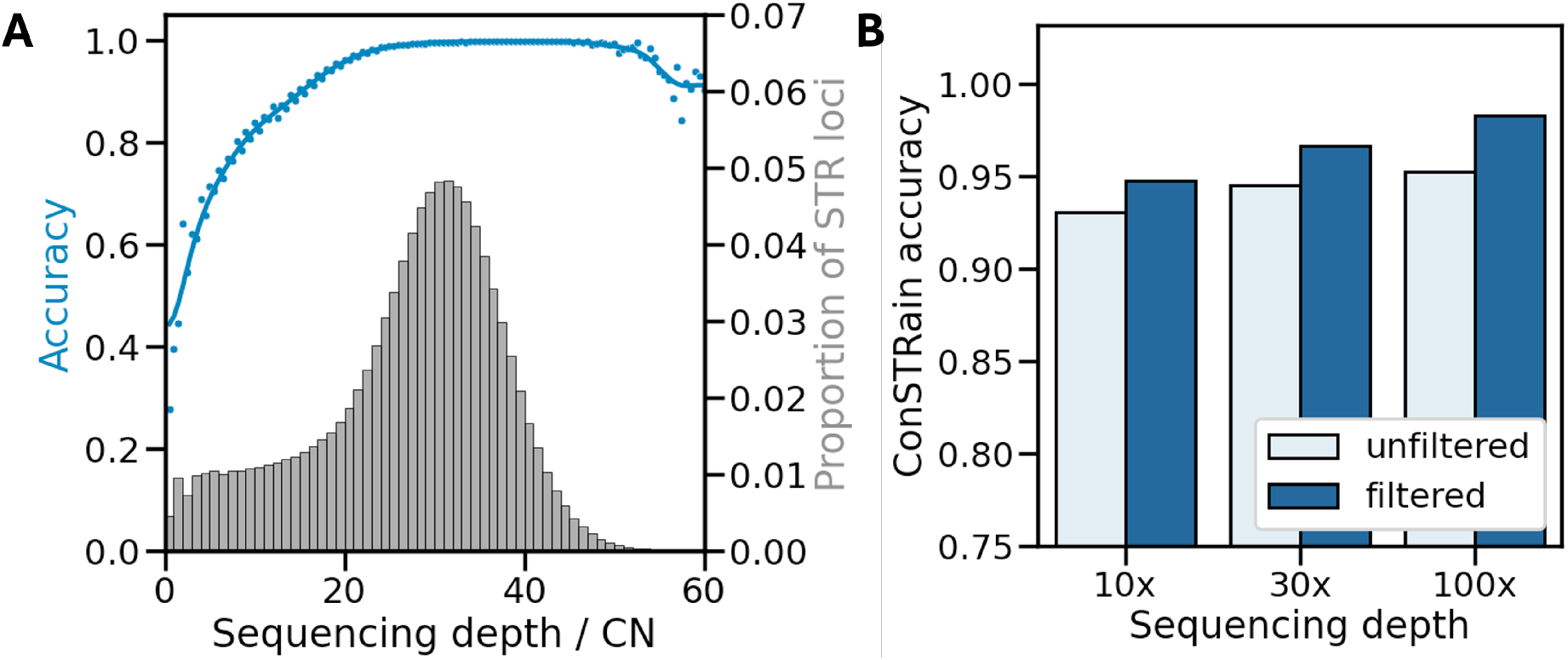
ConSTRain performance on Q100 benchmark. (**A**) Distribution of normalised sequencing depth observed by ConSTRain across 167114 repeat loci in the 100X HG002 WGS sample. The x-axis shows the sequencing depth normalised by the copy number of repeat loci. The left y-axis shows the accuracy of allele length calls (blue line and dots). The right y-axis shows the proportion of loci (grey histogram). Note: only normalised depth values between 0 and 60 are shown for visual clarity. (**B**) Accuracy of unfiltered and filtered ConSTRain STR allele length calls for 100X WGS of HG002, as well as for the same sample downsampled to 30X and 10X depth of coverage. Note: y-axis starts at 0.75.

**Fig. 3:**
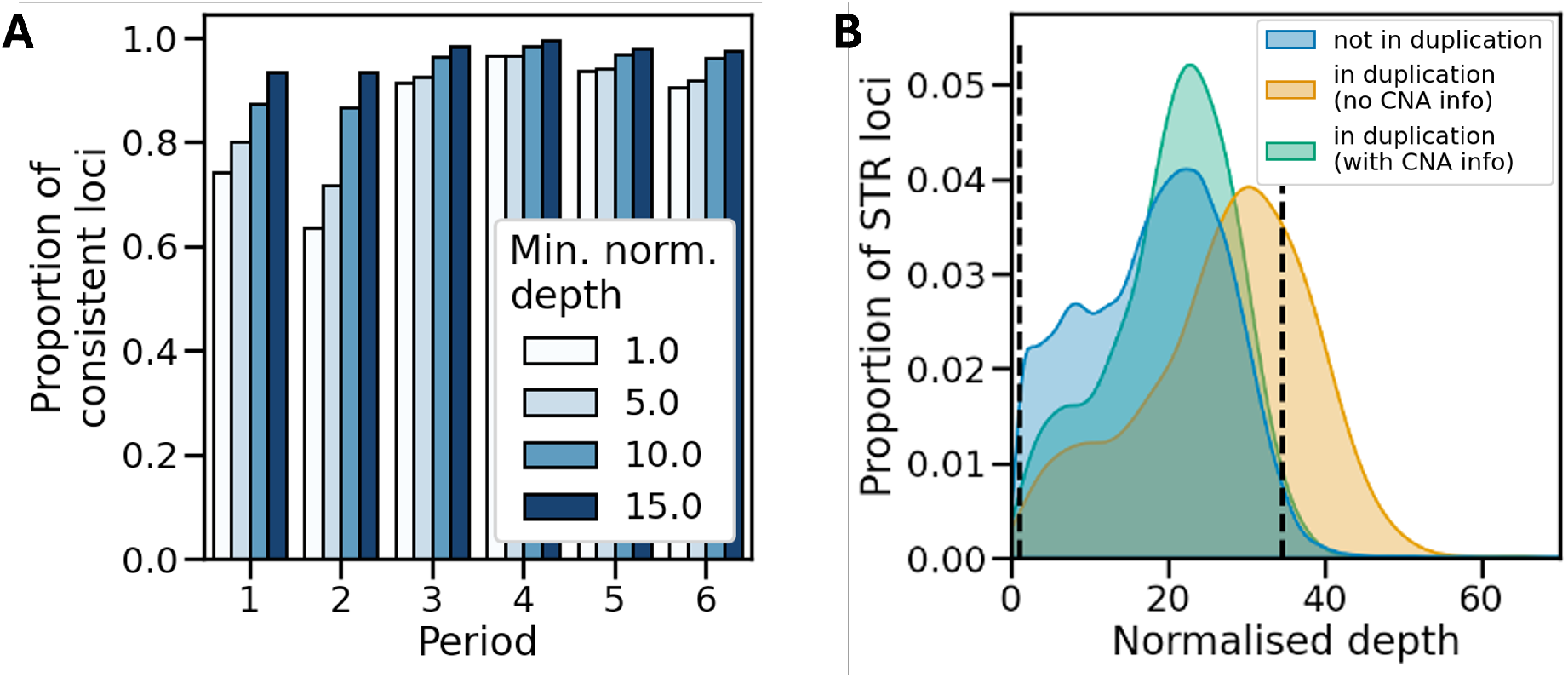
Genotyping STRs in a triploid *M. acuminata* sample with a large duplication on chr02. **(A)** Consistency of STR genotypes between the HiSeq1500 and NextSeq500 samples for different normalised depth filtering thresholds. X-axis: STR period, y-axis: proportion of loci for which the inferred genotype matched exactly between the two alignments. **(B)** Distributions of the depth of coverage for STR loci normalised by copy number for STRs in the alignment of combined HiSeq1500 and NextSeq500 reads. The blue distribution shows the normalised depths for loci not affected by CNAs. The orange distribution shows the normalised depth reported for loci in the chr02 duplication when CNA information was not provided to ConSTRain. The green distribution shows normalised depth for the loci in the chr02 duplication when CNA information was provided to ConSTRain. Vertical dashed lines indicate filtering bounds that exclude the 2.5% of loci with the highest and the 2.5% of loci with the lowest depth of coverage in the overall sample.

Next, we investigated if accuracy could be increased by filtering STR loci based on their normalised depth (see Methods). To this end, we generated the distribution of normalised read depths shown in Figure 2A. The distribution is left-skewed. Upon further investigation, we found that this was caused by mononucleotide repeats, whereas the distribution for repeats with higher periods followed a normal distribution (Supplementary Figure 2). Therefore, we decided to not consider mononucleotide repeats when defining our filter parameters, and instead used the distribution of loci with periods greater than one. In this distribution, we set bounds such that they excluded loci that fell in the lowest 2.5% and highest 2.5% of normalised depth values. These bounds were then used as parameters to rerun ConSTRain on the VCF (including mononucleotide repeats) previously created from the HG002 alignment. Additionally, we filtered out loci that overlapped known segmental duplications in the human genome. Together, this decreased the number of called loci to 1393426 (82.17% of the total), but increased the accuracy to 98.28% (Figure 2B, Supplementary Figure 3).

Having found that ConSTRain was able to accurately determine repeat allele lengths in a 100X short-read sequencing alignment, we wanted to see how it performed on samples with lower sequencing depths. To examine this, we downsampled the HG002 alignment to 30X and 10X depth of coverage. The accuracy of unfiltered allele length calls was 94.51% and 93.06% for the 30X and 10X alignments, respectively (Figure 2B). Importantly, normalised depth-based filtering of loci proved to be effective for the downsampled alignments as well. After filtering the genotyping accuracy rose to 96.65% and 94.75% for the 30X and 10X alignments, respectively (Figure 2B).

### ConSTRain’s accuracy is competitive with existing STR variant callers

Next, we sought to compare ConSTRain’s performance to that of other STR variant callers. We ran GangSTR and HipSTR on the 100X HG002 alignment using the same STR reference panel used for ConSTRain and again compared reported allele lengths to the ground truth haplotypes. For both tools, we analysed their unfiltered outputs, as well as the outputs filtered according to instructions in the respective tool’s documentation. The results are shown in Table 1. The accuracy of the filtered outputs of the three methods are very similar: ConSTRain had an accuracy of 98.28%, GangSTR 97.69%, and HipSTR 97.74%. A notable difference between the three methods was that HipSTR called substantially fewer loci (69.28% of the STR reference panel) than both ConSTRain (82.17%) and GangSTR (80.95%). Another major difference was that ConSTRain had a much lower runtime than the other two tools. When running single-threaded, ConSTRain was around 2.2 times as fast as GangSTR, and 1.8 times as fast as HipSTR. Moreover, ConSTRain supports running on multiple threads. With 32 threads, ConSTRain took 19 minutes and 31 seconds to genotype our reference panel of over 1.7 *×* 10^6^ loci in the 100X HG002 alignment, making it 45.8 times as fast as GangSTR and 36.9 times as fast as HipSTR in this benchmark.

**Table 1.**
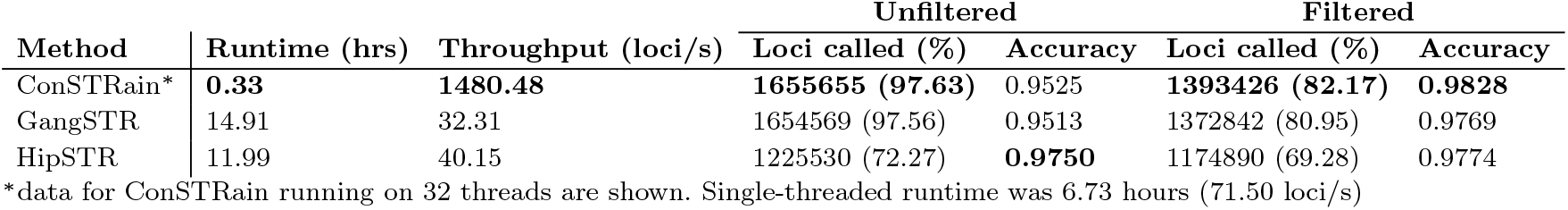
Results for ConSTRain, GangSTR, and HipSTR on the HG002 human benchmark. Data are shown before and after filtering the output of each tool. The total number of loci in the benchmark was 1695865. The percentage of this number that was called by each variant caller is shown in brackets in the ‘Loci called’ columns. The best value in each column is printed in bold.

### ConSTRain resolves STR genotypes in a simulated trisomy 21 sample

Having demonstrated that ConSTRain accurately recovers STR allele lengths from sequencing data of a diploid genome, we were curious to see how it performed at other copy numbers. We mimicked a trisomy 21 event by simulating short sequencing reads from three different assemblies of the human chromosome 21 and mapping them to GRCh38 (see Methods). We ran ConSTRain on the resulting alignment, specifying that the ploidy of chromosome 21 was three. After filtering, ConSTRain was able to estimate a genotype for 18241 out of the 21482 loci located on chr21 in our reference panel. At 13465 of these, the ground truth consisted of one distinct allele length (genotype *AAA*) across the three input assemblies. For 3923 and 853 loci two (genotype *AAB* ) and three (genotype *ABC* ) distinct allele lengths were present in the input haplotypes, respectively. Overall, ConSTRain reported the correct genotype for 98.39% of loci. Accuracy depended on the number of distinct alleles at a locus: ConSTRain reported the correct genotype for all loci with genotype *AAA*, 94.88% of loci with genotype *AAB*, and 92.76% of loci with genotype *ABC*. Similar to the HG002 benchmark, the majority of errors were because the distribution of generated allele lengths was not representative of the underlying genotype. This is possible because there was some stochasticity in the simulation of sequencing reads from reference haplotypes with regards to the depth of coverage along the input sequence. For example, ConSTRain reported an incorrect genotype for a mononucleotide repeat for which the genotype across the three haplotypes consisted of two alleles of length 19 and one of length 20. For this locus, 35 spanning reads with allele length 19 had been generated, and only three reads with allele length 20. Therefore, ConSTRain reported a genotype consisting of three alleles of length 19. Similar observations were made for other incorrectly called loci.

We were also interested to see what genotypes HipSTR and GangSTR would report in this setting. We ran both variant callers on the alignment of simulated reads without filtering and parsed their outputs. Both tools reported homozygous genotypes for all repeat loci with genotype *AAA*. For loci with genotype *AAB* they usually reported a genotype that was homozygous for the more abundant of the two alleles, missing the other allele length. In a subset of these loci (7.53% for GangSTR, 6.49% for HipSTR) a heterozygous *AB* genotype was reported. Finally, for the loci with genotype *ABC*, both tools almost always reported heterozygous genotypes containing two of the three alleles. Which of the three alleles at a locus was ignored seemed to be dictated by which allele was represented by the fewest reads.

### ConSTRain accounts for CNAs in a triploid *Musa acuminata*

Given that we could accurately resolve STR genotypes in simulated reads of a triploid chromosome, we were curious how ConSTRain performed on a real polyploid sample. We obtained WGS reads from a *M. acuminata* Dwarf Cavendish banana, which is a triploid, and mapped them to the DH-Pahang v4 reference genome [20]. This particular sample was reported to have a duplication of around 6 megabases on the long arm of chromosome 02 [13], making it an even more relevant test case for ConSTRain.

There was no ground truth available for STR genotypes in this analysis. However, there were two separate sequencing experiments performed for the same sample (see Methods). Ideally, STR genotypes reported by ConSTRain should be consistent between the two samples [27]. We tested for consistency by running ConSTRain on the NextSeq500 and HiSeq1500 alignments separately (including coordinates of the chr02 duplication), and comparing STR genotypes between the two outputs (Figure 3A). We considered only genotypes that were exactly the same between the two outputs to be consistent. Initially, we again set the minimum and maximum normalised depth values such that 2.5% of the lowest and 2.5% of the highest depth loci with periods > 1 were excluded from each sample. Even with these rather lenient filter parameters, genotype calls for STRs with periods > 2 were consistent at over 90% of loci (Figure 3A). Genotype calls were much less consistent for mononucleotide (74.29% of calls consistent) and dinucleotide (63.66% of calls consistent) repeats, however. Consistency between samples could be increased by raising the minimum normalised depth value, also reaching around 90% for mono- and dinucleotide repeats when using a threshold of 10. or 15. (Figure 3A).

Next, we genotyped our banana STR reference panel using the merged alignment (see Methods), specifying that three copies existed of each chromosome but without providing coordinates for the chr02 duplication. ConSTRain ran in 70 seconds on 16 threads and reported genotypes for 153167 out of 183345 STRs in the panel before filtering. Afterwards, we ran ConSTRain on the resulting VCF file, this time providing coordinates of the duplicated region (Supplementary Figure 4). We found that 2699 of the genotyped STR loci were located in the duplicated region. Figure 3B shows the distribution of normalised depths of coverage reported by ConSTRain across STRs. The normalised depth distribution for STRs in the duplicated region is shown separately — both before and after providing duplication coordinates to ConSTRain. The mean normalised depth for the non-amplified STRs was 18.84, while the mean normalised depth for the amplified STRs was 27.29. This difference in mean normalised depth is roughly in line with a tetraploid region being mistakenly analysed as triploid (theoretically, normalised depth at tetraploid loci should be 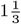 times that of the normalised depth in triploid loci in this case).

**Fig. 4:**
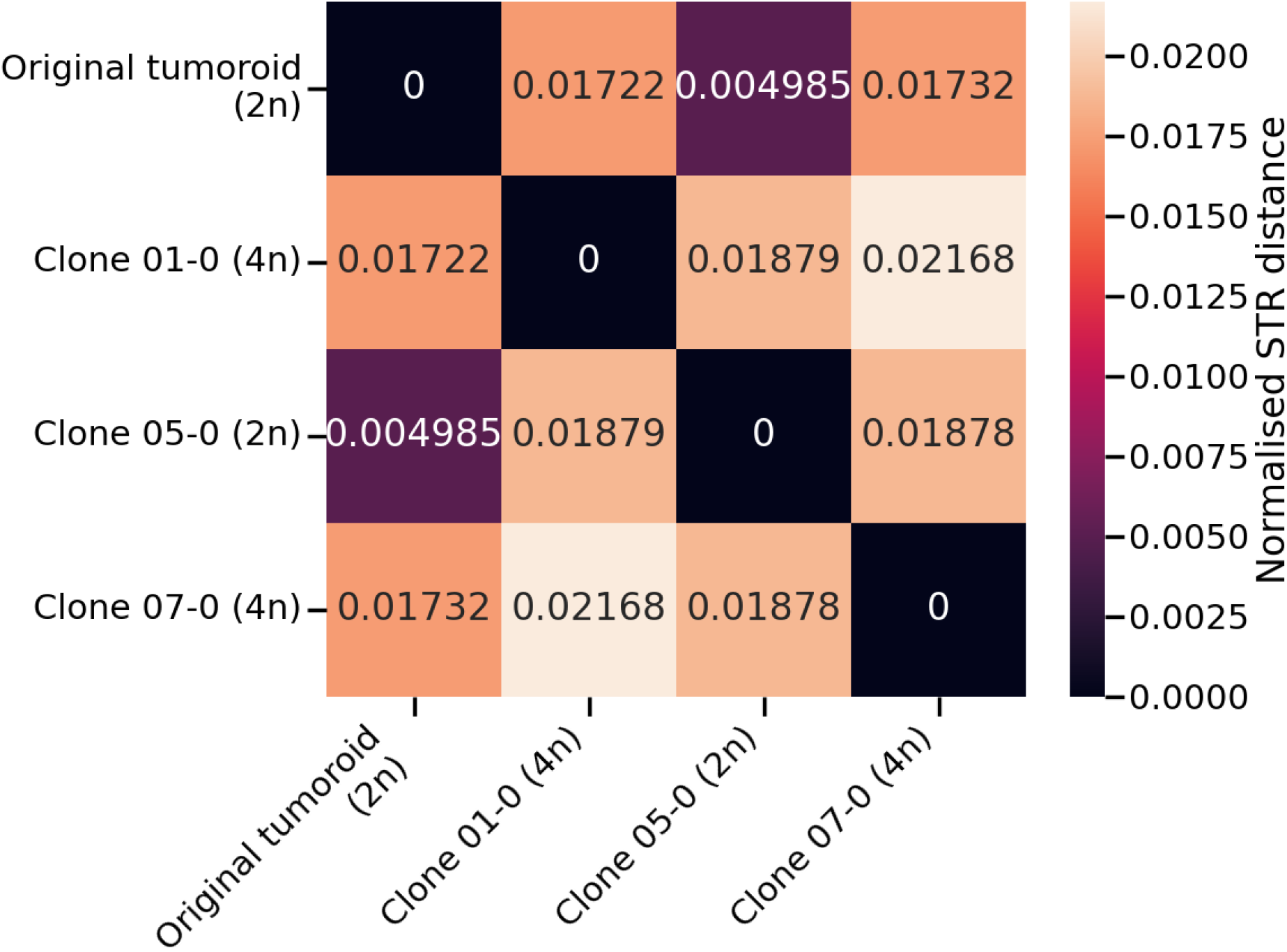
Pairwise STR-based distances between four samples stemming from the same patient-derived tumoroid. Each cell represents the comparison between two samples, with the colour and value of cells indicating the normalised distances between samples (average difference in allele length per locus).

Furthermore, the distribution of normalised depths reported for amplified STRs when the duplication coordinates were provided to ConSTRain largely overlapped the distribution for loci in the rest of the genome (Figure 3B). This highlights an additional benefit of setting filter parameters based on the normalised depth of loci observed in a sample: using bounds such that 2.5% of the lowest and 2.5% of the highest normalised depth loci with periods > 1 were excluded genome wide, 27.04% of STRs in the duplicated region were excluded when running ConSTRain without CNA information. When CNA information was provided to ConSTRain, however, only 3.44% of duplicated loci were excluded, since their normalised depth values were much more in line with the rest of the genome. This indicates that when CNA information is not available or is incorrect, a substantial portion of loci with incorrect copy number values may be excluded by filtering on normalised depth of coverage. This effect is expected to be even stronger when the difference between the annotated and true copy number of loci is larger.

After filtering, ConSTRain reported genotypes for 148532 STRs, 2612 of which were located in the duplicated region on chr02. At 55.15% of the triploid loci ConSTRain reported one distinct allele, 33.02% had two disctinct allele lengths, and 11.83% had three distinct alleles (Supplementary Figure 5). This distribution was very similar for loci located in the duplicated region on chr02 (Supplementary Figure 5). Interestingly, we also observed 31 loci with four distinct alleles in the duplicated region, 18 of which had a normalised depth of coverage *≥* 10. While this was only a small fraction of loci in the duplicated region, it suggests that some STR loci may have mutated after the duplication event.

### ConSTRain resolves STR genotypes in whole-genome duplicated colorectal cancer

Finally, we were curious to see whether ConSTRain can be used to study STRs in cancer sequencing data. To this end, we obtained four WGS samples that were derived from a microsatellite instable CRC tumoroid (see Methods) [14]. Of the four WGS samples, one was taken directly from the original tumoroid line. The other three samples represented clones 01-0, 05-0, and 07-0 that had been grown from single cells taken from the original tumoroid line, where clones 01-0 and 07-0 had undergone whole-genome duplications.

We ran ConSTRain on these four samples, providing CNA information each time. Next, we calculated STR-based pairwise distances between samples by comparing the STR genotypes (see Methods). Even though the four samples were all derived from the same tumoroid with only six weeks between the isolation of individual cells and the sequencing of the resulting clones, there were already some differences in STR genotypes between samples (Figure 4). The smallest distance was observed between the original tumoroid line and the diploid clone 05-0. The two tetraploid clones were more distinct from the original tumoroid line, with the STR-based distances between the original tumoroid line and both tetraploid clones being roughly similar. The largest pairwise distance we observed across this dataset was between the STR genotypes of the two tetraploid clones (Figure 4).

## Discussion

To the best of our knowledge, ConSTRain is the first STR variant caller that enables rapid and accurate analyses of microsatellites while accounting for copy number alterations and polyploidy.

On a benchmark of a euploid human genome ConSTRain reached a genotyping accuracy of 98.28%, which was competitive with state-of-the-art STR variant callers. ConSTRain was faster than the two other STR variant callers included in our analyses, even when running single threaded. ConSTRain’s runtime can be easily reduced even further by using multiple compute threads: for example, when running on 32 threads, ConSTRain genotyped over 1.7 *×* 10^6^ repeats in under 20 minutes from an alignment of 100X WGS reads. We showed that ConSTRain’s genotyping accuracy remained high for a simulated trisomy 21 event, even at loci with three distinct alleles. Then, we demonstrated ConSTRain’s ability to call STR genotypes from sequencing data of a polyploid *M. acuminata* Dwarf Cavendish banana while accounting for a large amplified region on chromosome 02. The final analysis we presented here focused on four WGS samples from an MSI CRC tumoroid. Interestingly, two of these samples had undergone whole-genome duplications and were tetraploid. This is uncharacteristic for microsatellite instable tumours, which are typically chromosomally stable [14]. When we determined STR-based pairwise distances between all four samples, we found that the comparison between the two tetraploid clones yielded the largest distance in this dataset. This could indicate that these two clones are derived from different lineages within the tumour, meaning that at least two separate whole-genome duplication events occurred. While a more in-depth analysis is needed to determine this for certain, we have shown that ConSTRain could be used for such a study.

As with any method, it is important to be aware of ConSTRain’s limitations. In our HG002 benchmark and trisomy 21 analyses, we observed two main sources of errors. First, for some STR loci the observed allele length distribution strongly deviated from the underlying genotype, with some alleles being over- or underrepresented in the distribution. This was the largest source of errors in the HG002 benchmark (79.65% of errors were of this type). We do not consider this an inherent failing of ConSTRain’s genotype estimation approach: it reported the most likely genotype based on the observed allele length distribution in each case. However, it does highlight that an incorrect inference will be made if the observed read distribution strongly deviates from the underlying genotype. This is an issue for variant calling in general, and not specific to our method.

The second source of errors in the HG002 benchmark was due to rare instances where STR loci had an insertion or deletion that did not consist of an addition or removal of one or more complete repeat units. Similar to other STR variant callers [6, 7], ConSTRain only considers in-phase indel mutations, and thus does not return the correct allele lengths at such loci. The fact that this was observed at only 4873 out of 1393426 repeats in the filtered 100X benchmark suggests that this heuristic is reasonable. This is an opportunity for future extensions or updates to ConSTRain, however. To address out-of-phase indels ConSTRain’s core genotype inference approach would not need to be updated, but it would mean that the full sequence in each read mapping to repeat loci needs to be resolved. This is likely to increase runtimes compared to the current implementation, although it is impossible to say in advance by how much.

Another potential source of issues is the use of external CNA information, which ConSTRain does not validate. If incorrect information is provided, it is likely to result in incorrect genotype calls. Our *M. acuminata* analysis suggests this may be mitigated by setting filtering parameters based on the distribution of normalised depth values observed across loci in a sample (Figure 3B). An alternative approach could be to estimate the copy number of repeat loci as part of the method itself. However, since it is a non-trivial task and many existing tools are available, we decided not to implement this functionality in ConSTRain at this point.

Recent years have seen an increased appreciation of the regulatory roles STR variants play in human physiology and disease. We believe ConSTRain unlocks the extension of such findings to other settings and organisms. By studying how STR variability interacts with other structural variants, we hope to learn more about the role these highly variable genomic elements play in the context of cancer. Furthermore, some of the crops upon which we rely most for our food production have polyploid genomes. By leveraging ConSTRain to analyse microsatellites in these species we may discover more about how their genomes and phenotypes are regulated.

## Data Availability

Source code and precompiled binaries for ConSTRain, as well as STR reference panels, are available on the ConSTRain GitHub page: https://github.com/acg-team/ConSTRain. Scripts and notebooks that were used to perform analyses and generate visualisations included in this manuscript are available in a separate GitHub repository: https://github.com/acg-team/ConSTRain-analyses/tree/main. HG002 Q100 haplotypes can be downloaded according to instructions on the T2T consortium’s HG002 Q100 GitHub page: https://github.com/marbl/HG002/tree/main. The aligned Illumina sequencing reads for the HG002 cell line are hosted by NCBI and can be downloaded from https://ftp-trace.ncbi.nlm.nih.gov/giab/ftp/data/AshkenazimTrio/HG002_NA24385_son/NIST_Illumina_2x250bps/novoalign_bams/. The *Musa acuminata* sequencing data are hosted by the European Nucleotide Archive under study accession number PRJEB33317. The CRC tumoroid sequencing data are hosted by the European Genome-phenome Archive under accession number EGAD50000000411.

