## Supplementary Figures for "Genotyping Short Tandem Repeats Across Copy Number Alterations, Aneuploidies, and Polyploid Organisms"

### Supplementary Figures for "Genotyping Short Tandem Repeats Across Copy Number Alterations, Aneuploidies, and Polyploids" by

**Max A. Verbiest, Elena Grassi, Andrea Bertotti, and Maria Anisimova**

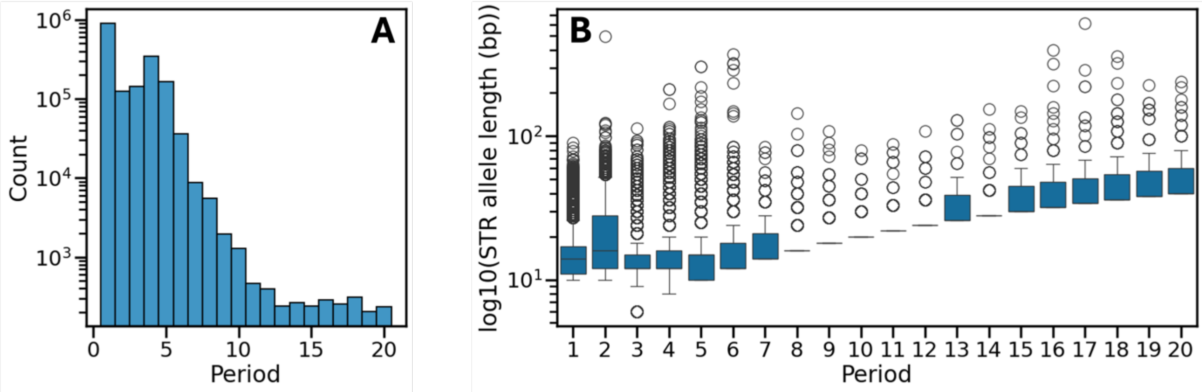


**Supplementary Figure 1.** Overview of repeat loci in the human STR panel used in this manuscript. **A)** Frequencies of different periods observed across the human STR panel. **B)** Boxplots showing the distribution of STR region lengths (in basepairs) for each STR period.

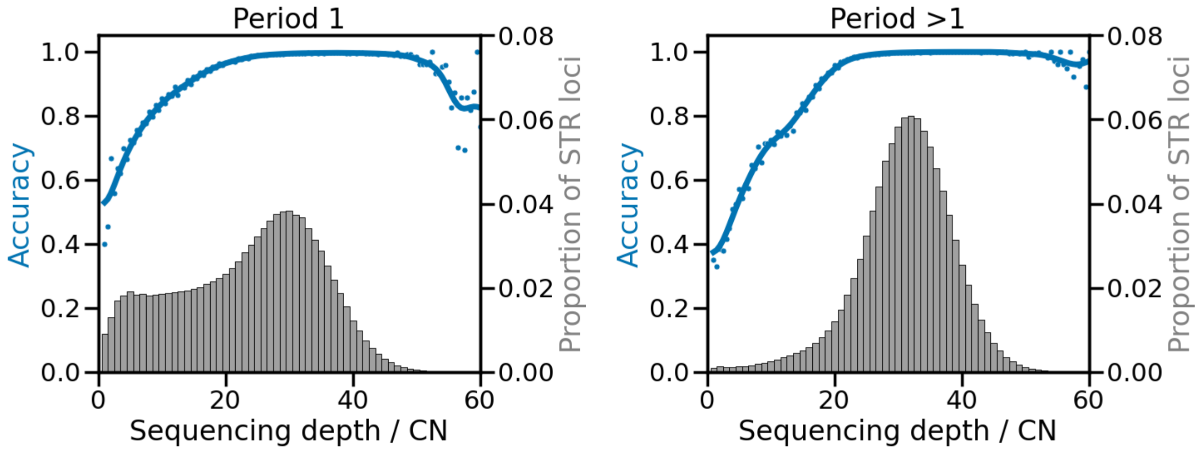


**Supplementary Figure 2.** Distribution of normalised sequencing depth observed by ConSTRain across repeat loci in HG002 WGS data. The left panel shows data for STRs with period 1. The right panel shows data for repeats with all other periods. X-axes show the sequencing depth normalised by the copy number of repeat loci. The left y-axes show the accuracy of allele length calls (blue line and dots). The right y-axes show the proportion of loci (grey histogram). Note: only normalised depth values between 0 and 60 are shown for visual clarity.

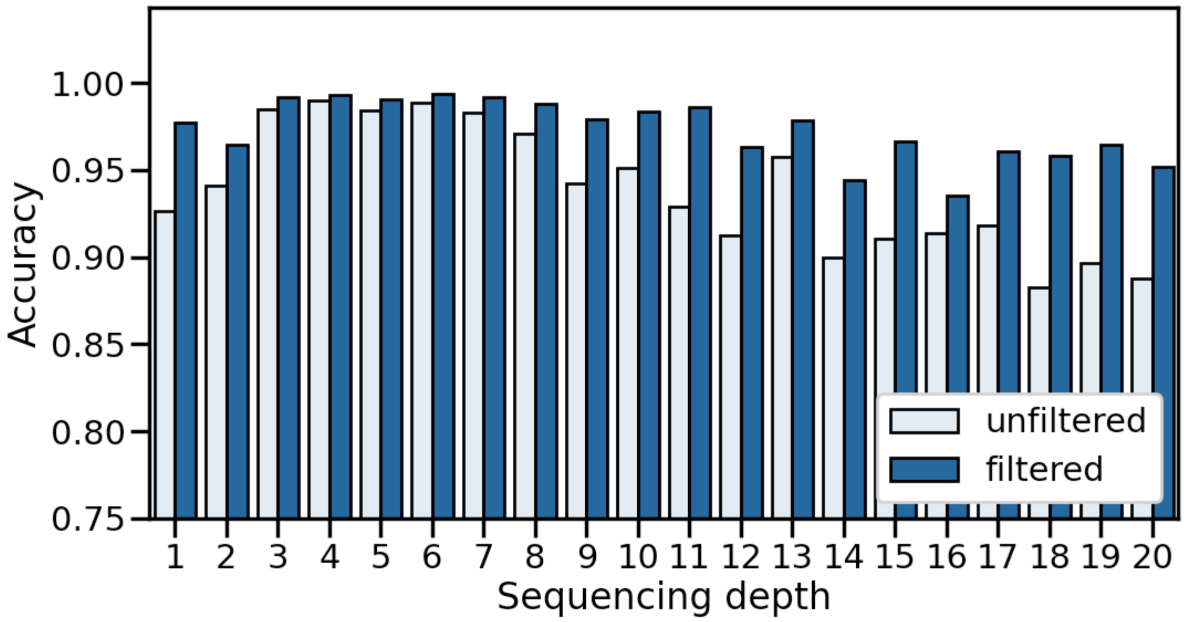


**Supplementary Figure 3.** Accuracy of ConSTRain allele length calls across repeat periods. Accuracy values were obtained by comparing genotypes reported by ConSTRain to the ground truth Q100 haplotypes generated by the T2T consortium. Accuracy values before and after filtering (removing loci in segmental duplications, removing loci with low depth of coverage) are shown. Please note: y-axis starts at 0.75.

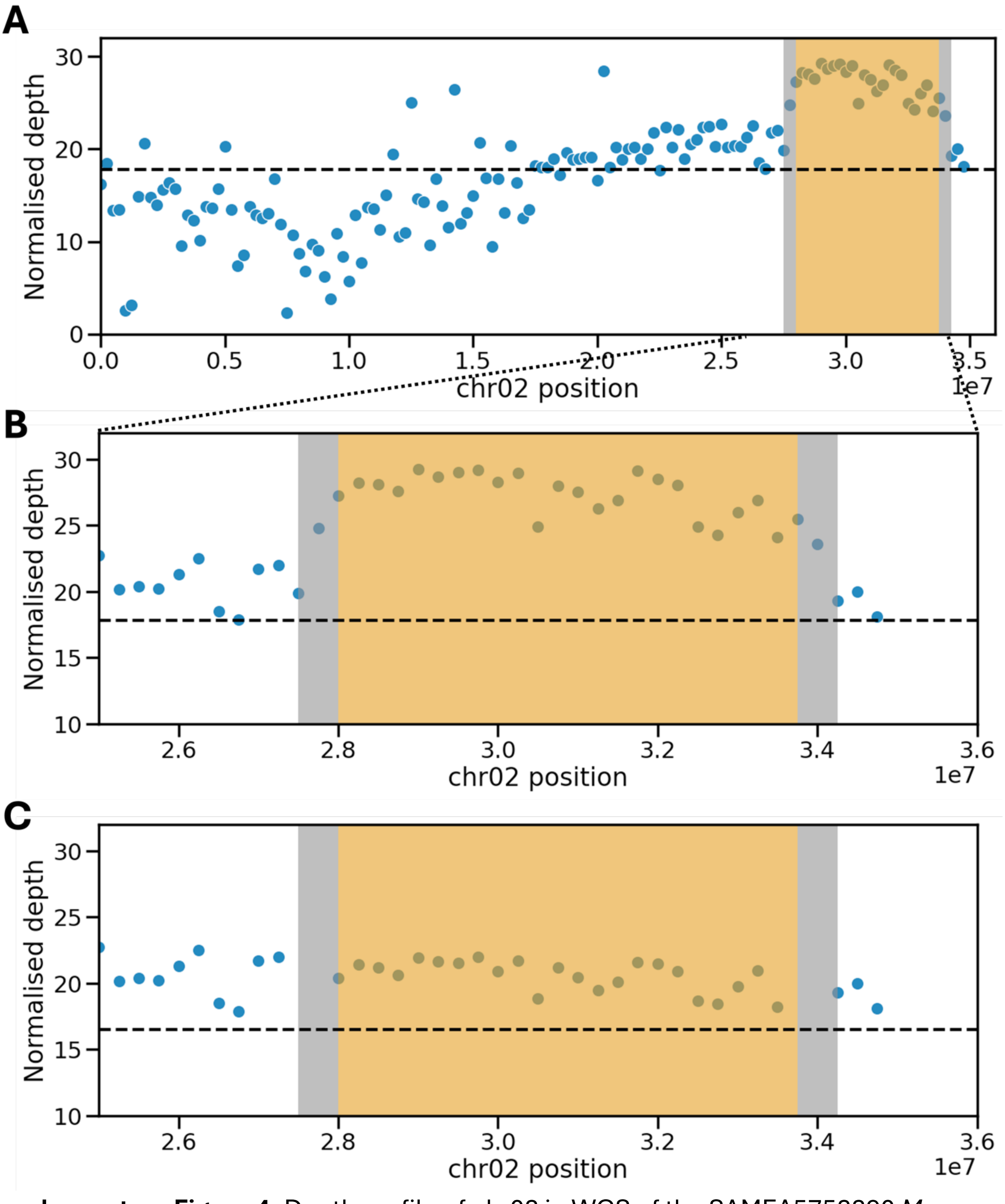


**Supplementary Figure 4.** Depth profile of chr02 in WGS of the SAMEA5752290 *Musa acuminata* sample. **A)** STRs on chr02 were assigned to bins of 2.5x$10^5$ base pairs based on their coordinates. The y-axis shows the normalised depth of coverage reported by ConSTRain for STR loci, averaged per bin. The horizontal dashed line indicates the average normalised depth of coverage for STR loci across the whole chromosome. Based on the depth profile, the previously reported duplication was visually identified (shaded in orange). **B)** Close-up of the amplified area. **C)** Same as B, but this time the y-axis shows normalised depth values reported by ConSTRain when coordinates of the duplicated region were provided. Since no accurate breakpoints for the amplification were available, STRs located within 500kb of the orange region were discarded (shaded in grey).

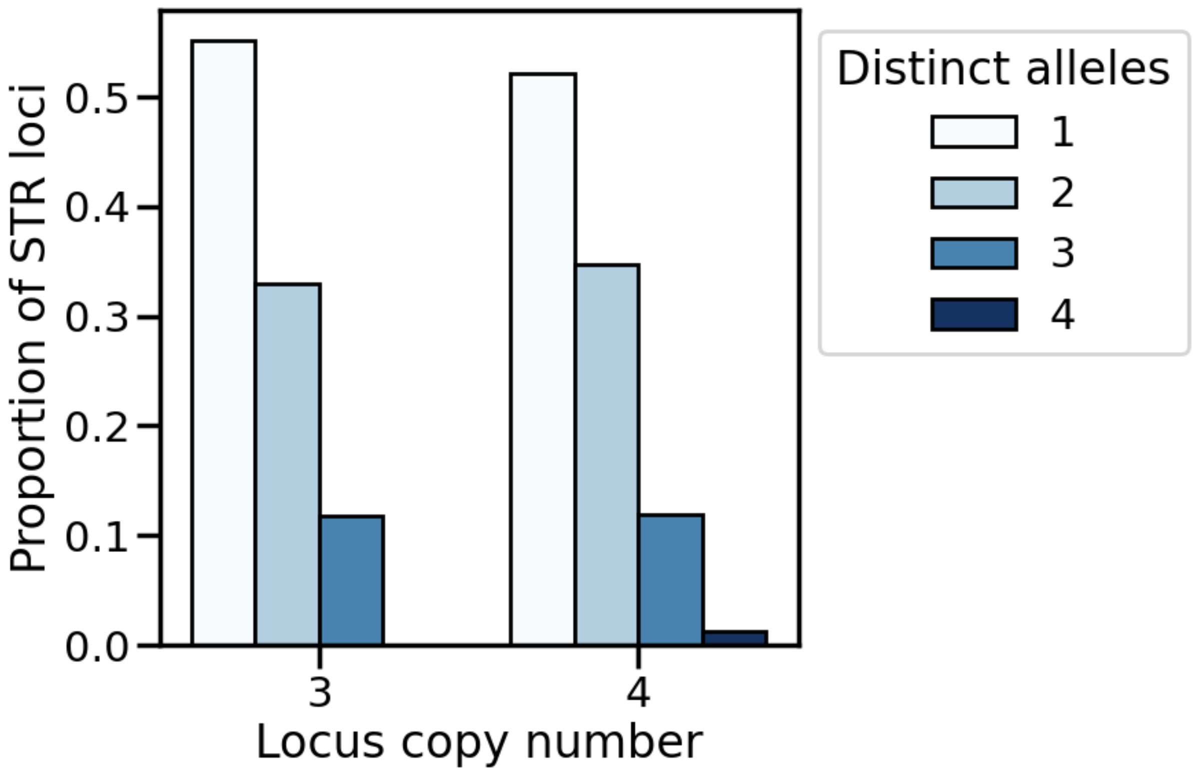


**Supplementary Figure 5.** The number of distinct alleles observed at STR loci in a *Musa acuminata* WGS sample. The x-axis shows the copy number for STR loci located in a duplication on chr02 (4) and the rest of the genome (3). Y-axis shows the proportion of loci for which a certain number of distinct alleles was reported by ConSTRain.
